# Why is usefulness rarely useful

**DOI:** 10.1101/2024.04.12.589314

**Authors:** Fangyi Wang, Mitchell J. Feldmann, Daniel E. Runcie

## Abstract

Mate selection plays an important role in breeding programs. The usefulness criterion was proposed as a criterion for mate selection, combining information on both the mean and standard deviation of the potential offspring, particularly in clonally propagated outbred species where large family sizes are possible. Predicting mean values of offspring of a cross is generally easier than predicting the standard deviation, especially in outbred species where the linkage of alleles is often unknown and phasing is required. In this study, we developed a method for estimating phasing accuracy from unphased genotype data on possible parental lines and evaluated whether the accuracy was sufficient to predict family standard deviations of possible crosses using a set of simulations spanning a wide range of genetic architectures and genotypes from a real strawberry breeding population. We find that despite highly accurate computational phasing, predicting family standard deviations and using predicted values of the usefulness criterion per possible cross confers little benefit relative to simply selecting parents based on predicted family means. Therefore even in this species, which is clonally propagated, outbred, and produces large families, we find the usefulness criterion unlikely to be useful.

## Introduction

Plant breeding programs aim to create genotypes with improved performance in one or more target traits. New genotypes are created by making crosses between existing genotypes, using recombination and Mendelian assortment to assemble sets of alleles that did not exist. These new genotypes can then be evaluated and released as commercial products or recycled to create new crosses. The success of a breeding program can be improved by evaluating genotypes more efficiently and making crosses that are more likely to generate better genotypes (Bernardo 2014a; Meuwissen *et al*. 2001; Heffner *et al*. 2009; Forster *et al*. 2015). In recent years, genomic data has been leveraged in many breeding programs to make both steps more effective. Genomic data can help make the evaluation of genotypes based on phenotypic data more precise by modeling the correlations between phenotypes and genotypes. Genomic data can also be used to select parents to be crossed by predicting their breeding value (BV), or the expected average performance of their offspring (Meuwissen *et al*. 2001; Heffner *et al*. 2009; Jannink *et al*. 2010). Both uses of genomic data are now commonly implemented in plant breeding programs.

Predicting the breeding value of potential parents is widely used to choose crossing partners. When the number of offspring per cross is large, the best crosses are the ones that will create the best possible offspring, not the crosses that will make the best average offspring. This situation is common in many plant systems because large numbers of seeds can be generated from the same cross. This result was observed nearly 50 years ago by Schnell and Utz (1975), who proposed the usefulness criterion (UC), or usefulness, to design optimal crosses (Zhong and Jannink 2007). Usefulness is defined as

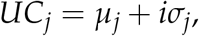

where *µ*_*j*_ is the average genetic value of offspring of cross *j, σ*_*j*_ is the standard deviation (SD) of genetic values among these offspring, and *i* is the selection intensity, the difference in mean phenotypic value after selection and before. The consequence of the usefulness is that crosses between parents that individually have good breeding values are not necessarily the best to make. For example, the upper tail of the distribution of genetic values of offspring from a cross with high variance (high *σ*^2^) may have higher values than the upper tail of the distribution of offspring from a family with higher mean but lower variance. Selection decisions based on usefulness have received little attention in animal breeding, possibly because the number of offspring per cross is very small, so the chance of creating offspring in the tails is low. However, several recent plant breeding studies have suggested that usefulness-based breeding might be successful, particularly in clonally propagated crops (Wolfe *et al*. 2021).

A simple method to predict the mean breeding value of a family is to use the average of the breeding value of the two parents. This will be accurate as long as most QTL operate additively. However, even under additive gene actions, predicting the variance within a family is more challenging as patterns of co-segregation of causal loci among genotypes also contribute to variance. For example, a cross between two genotypes with two coupling heterozygous loci would produce a greater variance in the next generation, compared to one with two repulsion heterozygous loci (Lynch and Walsh 1998; Zhong and Jannink 2007; Wolfe *et al*. 2021). Predicting co-segregation of causal loci from a cross of two outbred parents requires information on the phasing of alleles between all pairs of heterozygous loci in each parent and phased haplotypes are not directly measured by most genotyping platforms when individuals are heterozygous. Computational phasing using software like Beagle (Browning and Browning 2007; Browning *et al*. 2018) and FILLIN (Swarts *et al*. 2014) have been used in plant populations (Poland and Rife 2012; Scott *et al*. 2020). However, whether these methods are sufficiently accurate to predict family variance is unknown. Once phased haplotypes are known, QTL effect sizes need to be estimated in a training population, and then either family population variance (Zhong and Jannink 2007; Osthushenrich *et al*. 2017) or sample variance can be predicted (Zhong and Jannink 2007; Osthushenrich *et al*. 2017; Lehermeier *et al*. 2017; Bernardo 2021; Wolfe *et al*. 2021).

The benefit of choosing crosses based on their usefulness will likely vary among crops and breeding programs due to differences in genetic architecture, breeding system, and training population size. A recent study by Wolfe *et al*. (2021) evaluated the usefulness in a cassava breeding program with pre-phased parental individuals and empirical observations of cross-variance in 4 traits. They argued that both family mean and variance were important metrics for mate selection. In their results, the family mean and usefulness criterion were correlated and similar selections were made using either criterion. This was one of the first applications of usefulness in an outbred, clonally propagated species using whole-genome marker data and genomic prediction models of marker effects. Our work focuses on strawberry, another outbred, clonally propagated species, at the University of California Davis (UCD) strawberry breeding program. This breeding program is currently investing in infrastructure to implement genomics data in breeding decisions (Pincot *et al*. 2022; Jiménez *et al*. 2023; Hardigan *et al*. 2023; Feldmann *et al*. 2023; Knapp *et al*. 2024) and has detailed records of the pedigrees of current varieties which help evaluate the success of the statistical models necessary for predicting usefulness (Pincot *et al*. 2021; Knapp *et al*. 2023; Feldmann *et al*. 2024) as detailed below.

Our study builds on the work of Wolfe *et al*. (2021) in three key ways. First, we develop and apply a method to measure the accuracy of computationally phased parental haplotypes which is necessary for accurately predicting usefulness. Second, our study uses a different reference population with a different pedigree topology and a different marker density. Therefore, our results test whether their conclusions are robust in other systems. Finally, we use numerical simulations of genetic architectures so that we can compare computationally predicted usefulness values to the actual (simulated) values without measurement error. We repeat these simulations across a range of additive genetic architectures from oligogenic to polygenic and from low to high heritability so that our conclusions can generalize to many possible target traits. Overall, our results show that despite highly accurate computational phasing, the conditions under which selection decisions based on usefulness could be useful are relatively rare. This is consistent with earlier results from Bernardo (2021) who studied predicting genetic variance within biparental populations created from inbred parents and concluded that there was little benefit to modeling within-family genetic variance.

## Methods and Materials

### Genetic Data and Genotyping Error Estimation

We used genetic and pedigree data from a population of 1035 strawberry genotypes at 37,441 SNPs. To access genotyping error rates and pedigree accuracy, we identified 417 family trios of two parents and one of their offspring based on the pedigree. These 417 offspring were not parents of any other genotypes in the pedigree. The remaining genotypes in the population (618) included the parents of the 417 offspring and genotypes that did not produce any offspring. Within each family trio, we examined SNP markers for which both parents were homozygous. We compared the parental and offspring genotypes at those markers and counted the number of mismatches between parents and the offspring. The genotyping error was calculated as the number of mismatched markers divided by the total number of homozygous markers compared per trio. All of the family trios had a genotyping error rate of <5%.

### Phasing

To simulate the offspring of a cross between outbred parents, the linkage between heterozygous alleles within each parent must be known. Assigning heterozygous genotype calls to haplotypes is called phasing. Beagle 5.0 (Browning and Browning 2007; Browning *et al*. 2018) was used, with version *05May22*.*33a* and seed 0, to phase the SNP data for all genotypes and to impute any missing genotype calls.

### Simulation of offspring

We randomly selected 520 unique pairs of strawberry genotypes as parents for the crossing simulation. We excluded 2,500 of the 37,411 markers due to their unknown map positions. For each pair of parents, we used the *Hypred* R package (Technow 2011) to simulate crossover events and produce gametes, and then the gametes were randomly paired to produce 200 offspring. Crossover events were simulated based on a count-location model. The number of crossover events (on a single chromosome) followed a Poisson distribution with the parameter *λ* = *L*, with *L* the length of the chromosome in Morgans. The locations of crossover events were sampled from a uniform distribution 𝒰 (0, *L*).

### Estimation of imputation and phasing error

Before calculating the mean and the variance of the simulated offspring, we measured the accuracy of phasing and imputation. We re-ran Beagle using the parents of the 417 trios, excluding all offspring. We estimated the imputation error using the same method of estimating the genotyping error as above. We compared genotypes of offspring to their parents (computationally phased) at pairs of markers for which at least one parent was heterozygous at both markers. We measured the percentage of offspring genotypes that were inconsistent with the predicted parental haplotypes for markers, binned by the recombination distance between markers.

We compared these rates of pairwise genotype inconsistencies to the expected rates for similar recombination distances by simulating phasing errors in the haplotypes of each parent and comparing to the simulated offspring genotypes using the “correct” parent haplotypes. We estimated the rate of offspring genotype inconsistency with the “incorrect” parental haplotypes using the method above. We defined 6 rates of phasing error per cM distance *p ∈* (0.005, 0.01, 0.025, 0.05, 0.1, 0.15) as the probability of a phasing error between two SNPs 1 cM apart. Phasing errors were simulated in the same fashion as crossover events as described above. The number of phasing errors per chromosome was sampled from a Poisson distribution with the parameter *λ* = *p ∗ L*. The locations of phasing errors were sampled from a uniform distribution.

### Simulation of breeding value and phenotypes

We simulated genetic values assuming a sparse additive genetic architecture. We created 30 scenarios spanning all combinations of six numbers of QTL (4, 16, 64, 252, 524, 1024) randomly selected from the sequenced SNP markers with effect sizes sampled from a standard normal distribution *N*(0, 1) and five levels of heritability *h*2 (0.1, 0.3, 0.5, 0.7, 0.9). We defined the breeding value (BV) of each genotype as the summation of the QTL effect sizes multiplied by the number of copies of the alternate allele, and simulated phenotypic values by adding random noise sampled from a normal distribution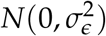, where 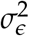 was scaled based on levels of heritability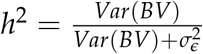. For each combination of heritability and QTL number,we ran 20 independent simulations, sampling different SNPs as QTL and with different effect sizes and environmental errors in each simulation. We calculated breeding values for both the real 1035 genotypes and simulated offspring, but the phenotypes were only simulated for the real 1035 genotypes.

### Prediction of family means and variances of breeding value, and estimation of prediction accuracy

RR-BLUP and BayesC models were fitted to the simulated phenotypes and the SNP marker data, excluding the markers that were selected as QTL, on the 1035 genotypes (Endelman 2011; Pérez and de los Campos 2014; Team 2020). Predicted breeding values were calculated for each offspring by plugging in offspring genotypes into the fitted models. We calculated the breeding value mean and breeding value standard deviation based on both the true and predicted breeding value of offspring with each family. We calculated the prediction accuracy as the Pearson correlation between the predicted values and the true values of each statistic (*cor*(*predicted*_*value, true*_*value*)).

### Calculation of selection intensity

The usefulness criterion is defined as a function of selection intensity *i*. To compare true and predicted usefulness values we selected four *i* spanning the range from weak to strong selection: 1.40, 1.75, 2.06, and 2.42, corresponding to selecting 20, 10, 5, or 2 percent of genotypes from populations with normally distributed phenotypes, using the equation:

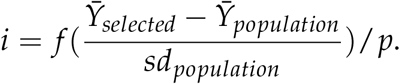

Here, *f* (*·*) is the probability density function of the distribution of the population, and we assume a normal distribution, and 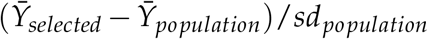 represents how many population standard deviations the selected group mean is away from the population mean. *p* is the proportion of genotypes selected.

## Results and Discussion

### Computational phasing is highly accurate in this strawberry population

We compared genotypes at pairs of markers at different genetic distances between unphased offspring and phased parents. To account for recombination, we binned marker pairs by genetic distances (in cM). Rates of inconsistencies increased with recombination distance (Figure 1), as expected. We then compared the observed rates of genotype inconsistency with expected rates if parents were correctly phased, or if phasing errors were introduced randomly within each parent with different frequencies. For marker pairs greater than 10 cM apart, our observed rate of genotype inconsistencies was close to the expected rate if parental phasing was perfect. However, for closer marker pairs, simulated inconsistency rates converged towards zero even when parental phasing was imperfect, but our observed inconsistency rate never dropped below *∼* 2.5%, perhaps due to a low amount of genotyping error or incorrect parent-offspring trios. These results are consistent with the computationally inferred phasing of markers being highly accurate in this population.

**Figure 1.**
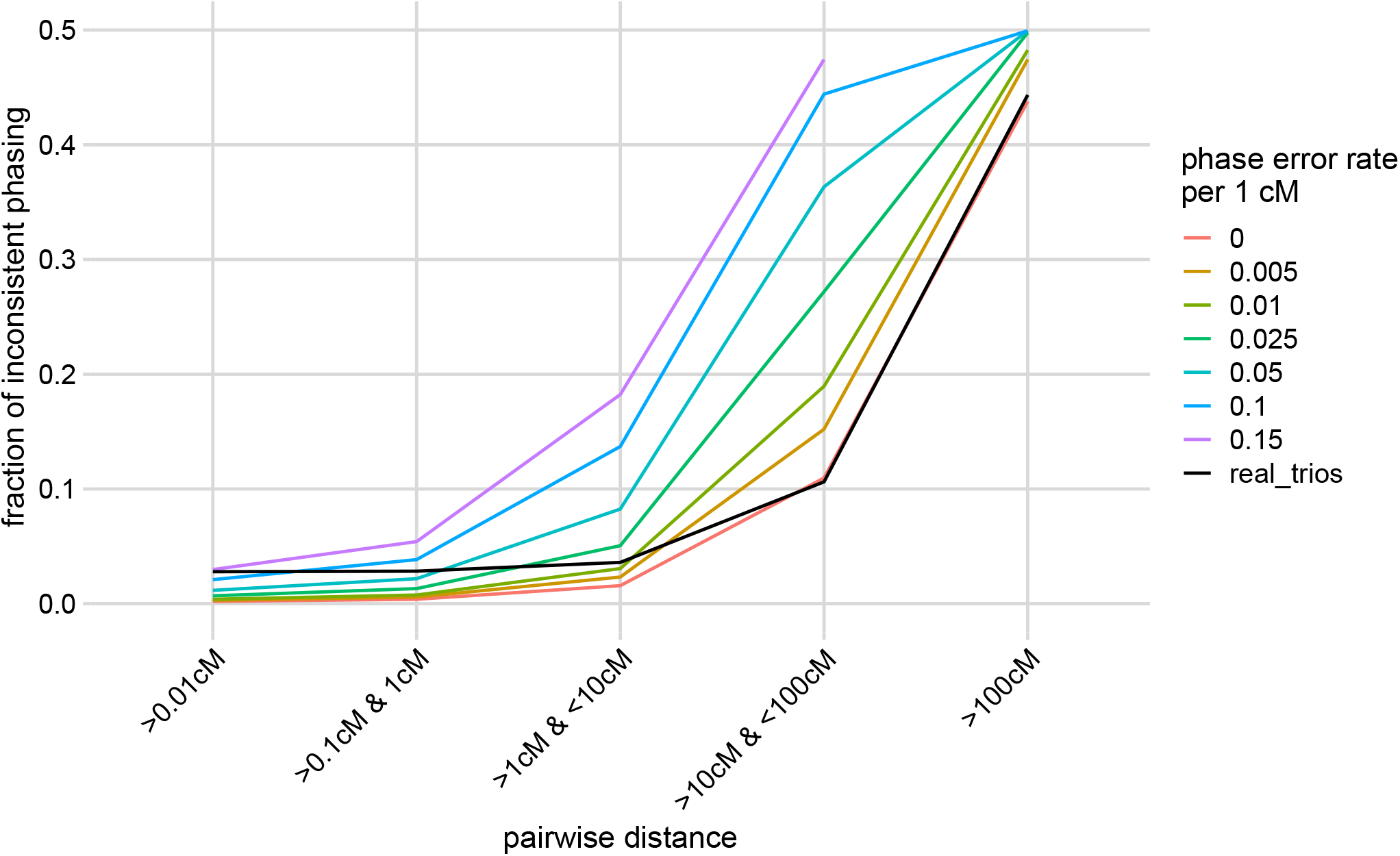
Rates of inconsistent parent-offspring genotypes at pairs of markers as a function of genetic distance. Genotypes at pairs of markers where both were heterozygous in at least one parent were compared between parents and offspring across the 417 family trios. The black line shows the fraction of marker pairs in each genetic distance bin inconsistent between offspring and parents. Colored lines show simulated inconsistency rates for the same marker pairs after introducing haplotype switches at the specified rates per cM in each parent.

### Prediction accuracy of family breeding value mean and standard deviation varied as heritability and number of QTL changed

Given the highly accurate computational phasing of parental haplotypes in this population (Figure 1), we used the haplotypes to simulate families of 200 offspring for 520 crosses between randomly paired genotypes and projected genotypes of causal alleles onto each simulated offspring. We then trained genomic prediction models (BayesC and RR-BLUP) on simulated parental phenotypes with varying levels of heritability (*h*2) and used the trained models to predict the genetic values of the offspring (Endelman 2011; Pérez and de los Campos 2014). We calculated the mean and standard deviation of these predicted genetic values and measured the accuracy of these predictions using Pearson’s correlation with the true values.

Using BayesC as a genomic prediction model, the average prediction accuracy of family means ranged between 60% and 90% (Figure 2a). We achieved moderate accuracy in predicting family standard deviations, averaging between 25% and 80% across scenarios (Figure 2b, red) Results using RR-BLUP as a prediction model were similar, but accuracy was slightly lower when heritability was high and the number of causal loci very low (supplementary). The prediction accuracies of family breeding values mean and standard deviations both increased as the heritability increased, as expected. When the number of QTL increased, the prediction accuracy of mean stayed the same (Figure 2a), but that of standard deviation decreased (Figure 2b, red). This might be due to less accurate estimates of individual QTL effects when there are more QTL.

**Figure 2.**
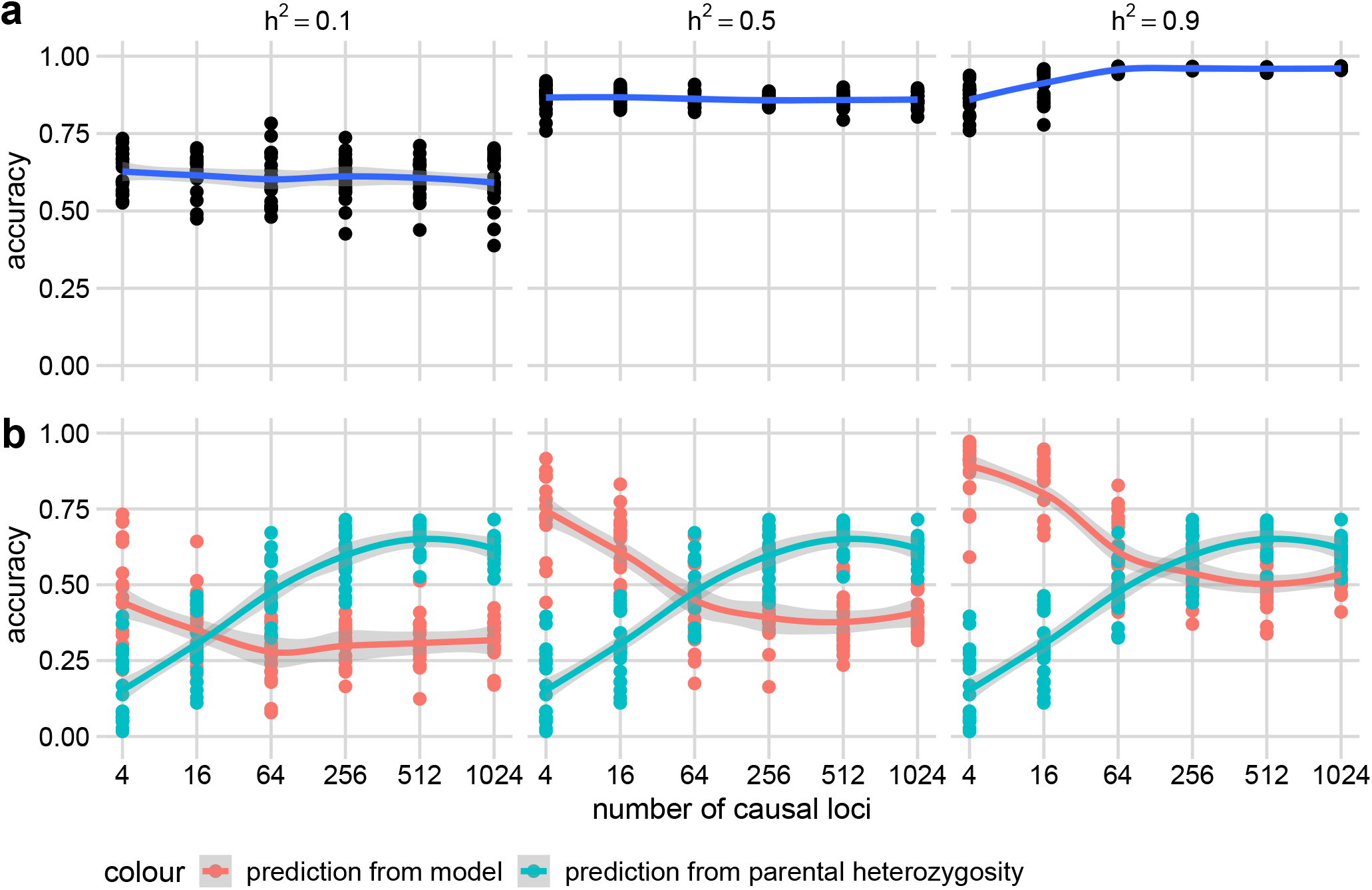
Prediction accuracies of the means (a) and standard deviations (b) of breeding values across simulated families as a function of heritability (*h*^2^) and the number of causal loci. Panels show Pearson’s correlations (accuracy, y-axis) between the true family means of breeding values (a) or true family standard deviations of breeding values (b, red) based on a BayesC genomic prediction model trained on parental phenotypes simulated with different heritabilities (*h*2) and different number of causal loci (x-axis). 20 simulations were run at each combination of *h*2 and numbers of causal loci, with different causal loci randomly sampled from the available SNP genotypes in each simulation. As a comparison, we also calculated the correlation between the true standard deviation of breeding values in each family and the average number of heterozygous markers of the two parents of that family calculated across all non-causal SNP markers in each simulation (b, blue). Points show measured accuracies in each simulation. Curves are calculated with the ‘geom_smooth()’ function and show smoothed estimates of the standard error of the mean accuracy across simulations.

Even at low heritability and high numbers of causal loci, prediction accuracies of standard deviation did not decline to zero. This is likely because parents heterozygous for more loci also tend to be heterozygous for more causal loci. Only heterozygous causal loci contribute to the variance of breeding values among offspring in our simulations because the genetic architecture is purely additive. The average parental heterozygosity of markers was more correlated with the standard deviation of offspring breeding values than the BayesC-based predicted standard deviations in most of our simulations (Figure 2b, blue), particularly when the number of causal loci was high. Mean heterozygosity information may have been indirectly captured by the genomic prediction models to accurately predict standard deviation. With a small number of QTL, the predicted family standard deviations from our genomic prediction models were more accurate than predictions of standard deviation from parental heterozygosity. In contrast, with a large number of QTL, the prediction of standard deviation from parental heterozygosity was more accurate. Unfortunately, despite its high correlation with family standard deviations, it is not straightforward to use parental heterozygosity directly to predict family usefulness because heterozygosity and family standard deviations have different units and scaling factors between trait units and heterozygosity units are unknown. In practice, real families with large enough numbers of genotypes are needed to estimate the regressions of family standard deviations and parental heterozygosity before this statistic could be used to estimate usefulness.

### Family usefulness can be accurately predicted with family mean

With the predicted family mean and standard deviation, we were able to predict the usefulness of each family for a defined selection intensity *i* using the equation: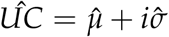. We calculated the accuracy of the estimated usefulness as Pearson’s correlation with the true usefulness (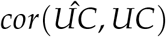) and compared this with the accuracy of predicting the usefulness using only the predicted family mean 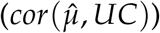 of each family. As heritability increased, the accuracy of both predictions increased (Figure 3). At low heritability, the prediction accuracies were relatively insensitive to the number of QTL. However, at high heritability, the prediction accuracy increased as the number of QTL increased. Predicting usefulness only achieved higher accuracies than predictions based on predicted family means when heritability was high and there were small numbers of QTL (Figure 3), the same conditions under which we could most accurately predict the family standard deviation (Figure 2b, red).

**Figure 3.**
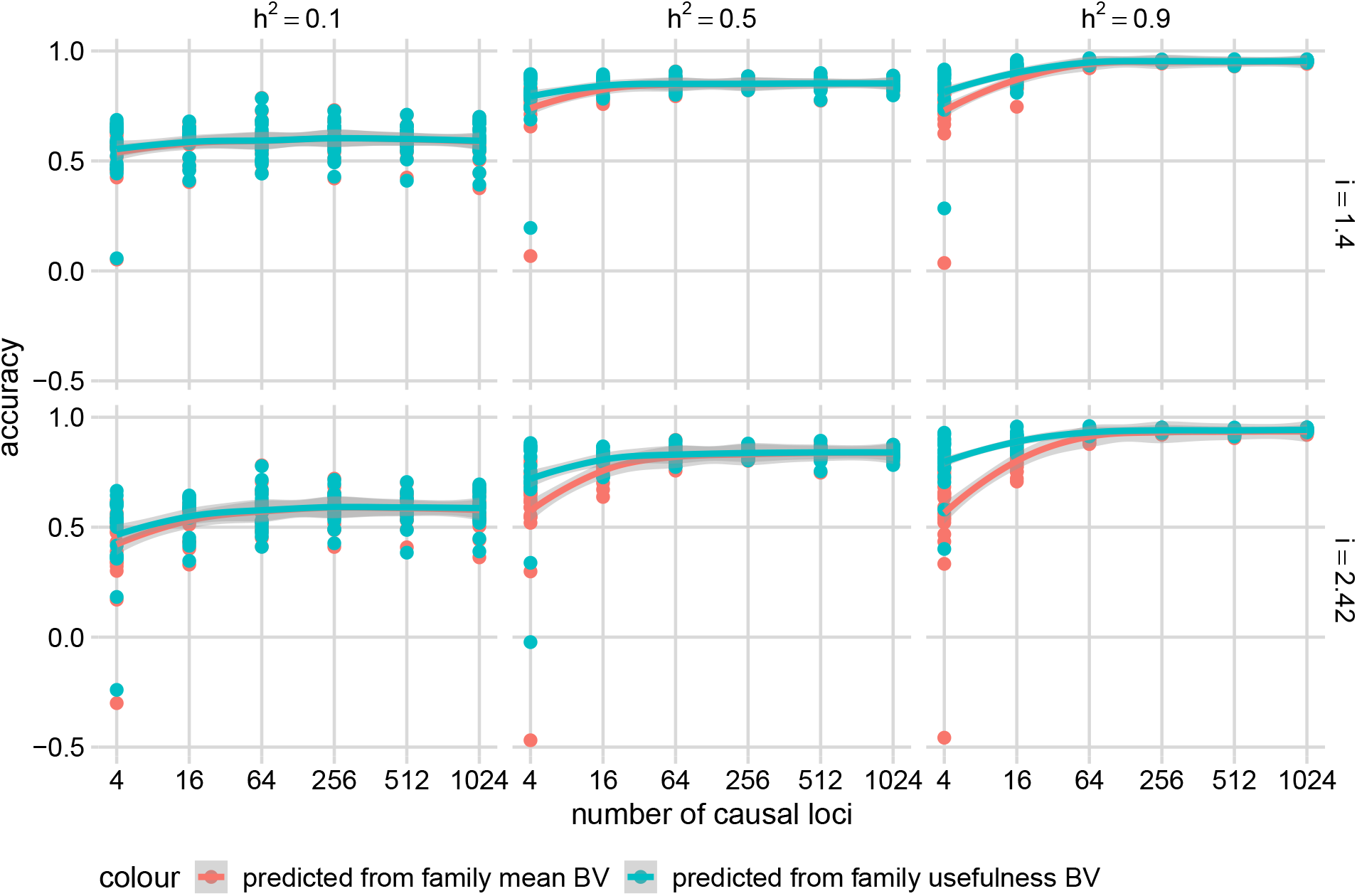
Prediction accuracies of the usefulness across the simulated families using a BayesC genomic prediction model. Panels show Pearson’s correlations (accuracy, y-axis) between the true family usefulness and predicted family mean (red) or predicted family usefulness (blue) based on a BayesC genomic prediction model trained on parental phenotypes simulated with different heritabilities (*h*2) and different numbers of causal loci (x-axis), across different selection intensity (*i*). 20 simulations were run at each combination of *h*2 and number of causal loci. Points show measured accuracies in each simulation across each *i*. Curves are calculated with the ‘geom_smooth()’ function and show smoothed estimates of the standard error of the mean accuracy across simulations.

Given the moderate accuracy at predicting both the standard deviation and mean of each family, we were surprised that direct predictions of usefulness were not much better than predictions based on predicted family means alone. One explanation for this observation is that the true values of family mean and family usefulness are highly correlated, especially when the number of causal loci is large (Figure 4a). The reason for this high correlation is that the variance among families in the quantity *iσ* (selection intensity multiplied by standard deviation) is very small compared to the variance among families in mean (*µ*) (Figure 4b), so the first term in the usefulness dominates.

**Figure 4.**
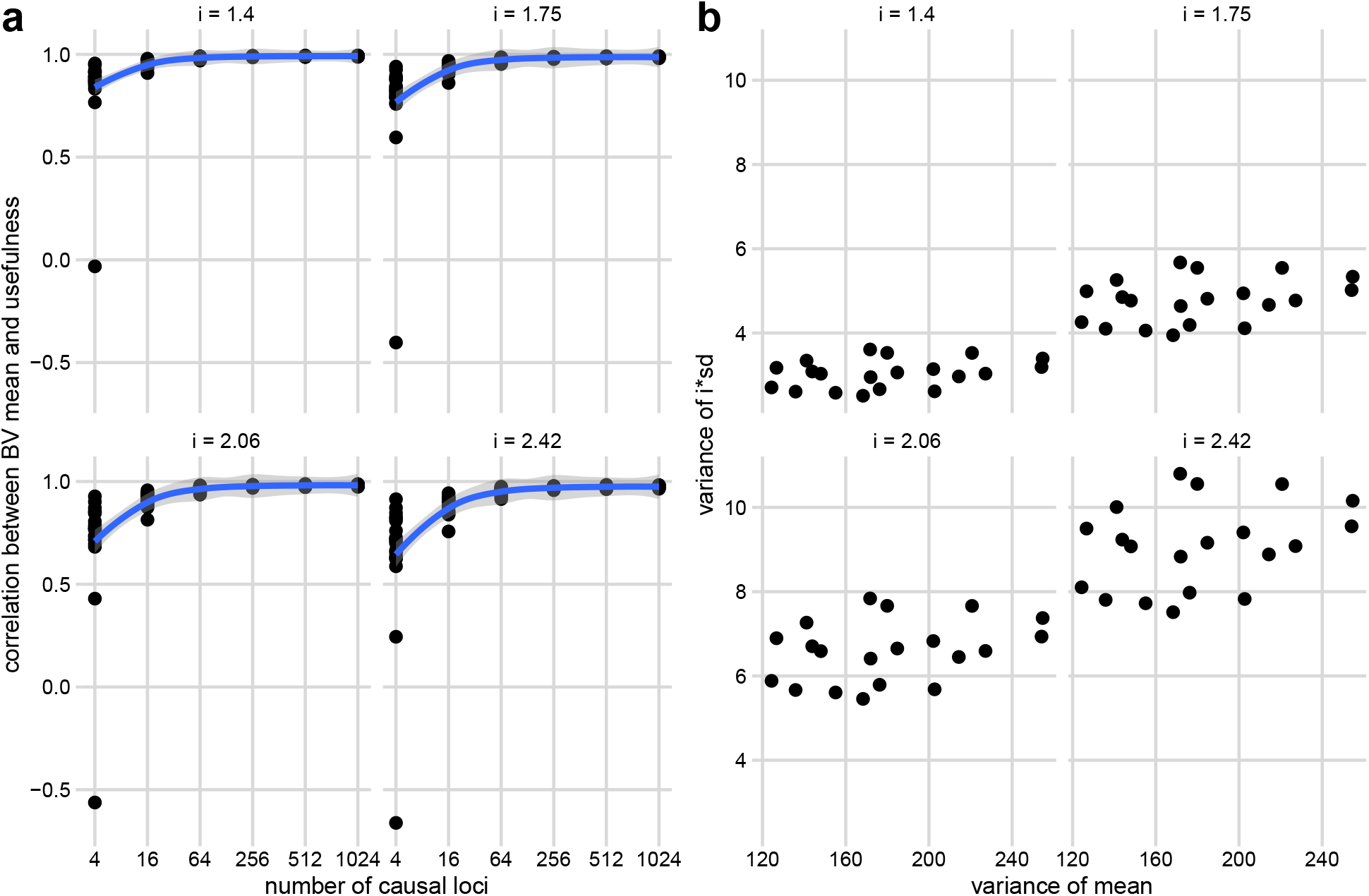
True family breeding value means and true family usefulness values are highly correlated. (a) Pearson’s correlation (y-axis) between the true family means of breeding values and true family usefulness across different selection intensities (*i*) and the number of QTL (x-axis). 20 simulations were run at each number of causal loci. Points show measured correlations in each simulation across each *i*. Curves are calculated with the ‘geom_smooth()’ function and show smoothed estimates of the standard error of the mean accuracy across simulations. (b) Scatterplots of the variance of selection intensity multiplied by standard deviation (y-axis) against the variance of the means (x-axis) for simulations with 1024 QTL. 20 simulations were run, and each point represents one simulation.

### Family usefulness cannot be accurately predicted with family mean in all circumstances

Zhong and Jannink (2007) previously discussed the utility of predicting family standard deviation, noting that if crosses among all candidate lines are evaluated for usefulness criterion, the variance of means will dominate the variance in standard deviation (Figure 4b). However, if crosses are limited to only among lines first selected for breeding value, the variance in means among candidate crosses will be reduced and the relative importance of standard deviation for usefulness will increase. To explore this, we simulated crosses among only the best 20 genotypes (by true breeding value) across the span of all genetic architectures and compared the correlation between the family means and family usefulness (Figure 5). The correlation between the family mean of breeding values and family usefulness was lower across all numbers of causal loci and selection intensities for these new families, reaching values as low as 0 when selection intensities were strong and there were few QTL. However, when the number of QTL was greater than 250, the correlation remained greater than 0.75 even for the highest selection intensity. Additionally, in practice, the best 20 genotypes would not be known *a priori*, and if some lower-performing genotypes were selected, the variance in family means would increase.

**Figure 5.**
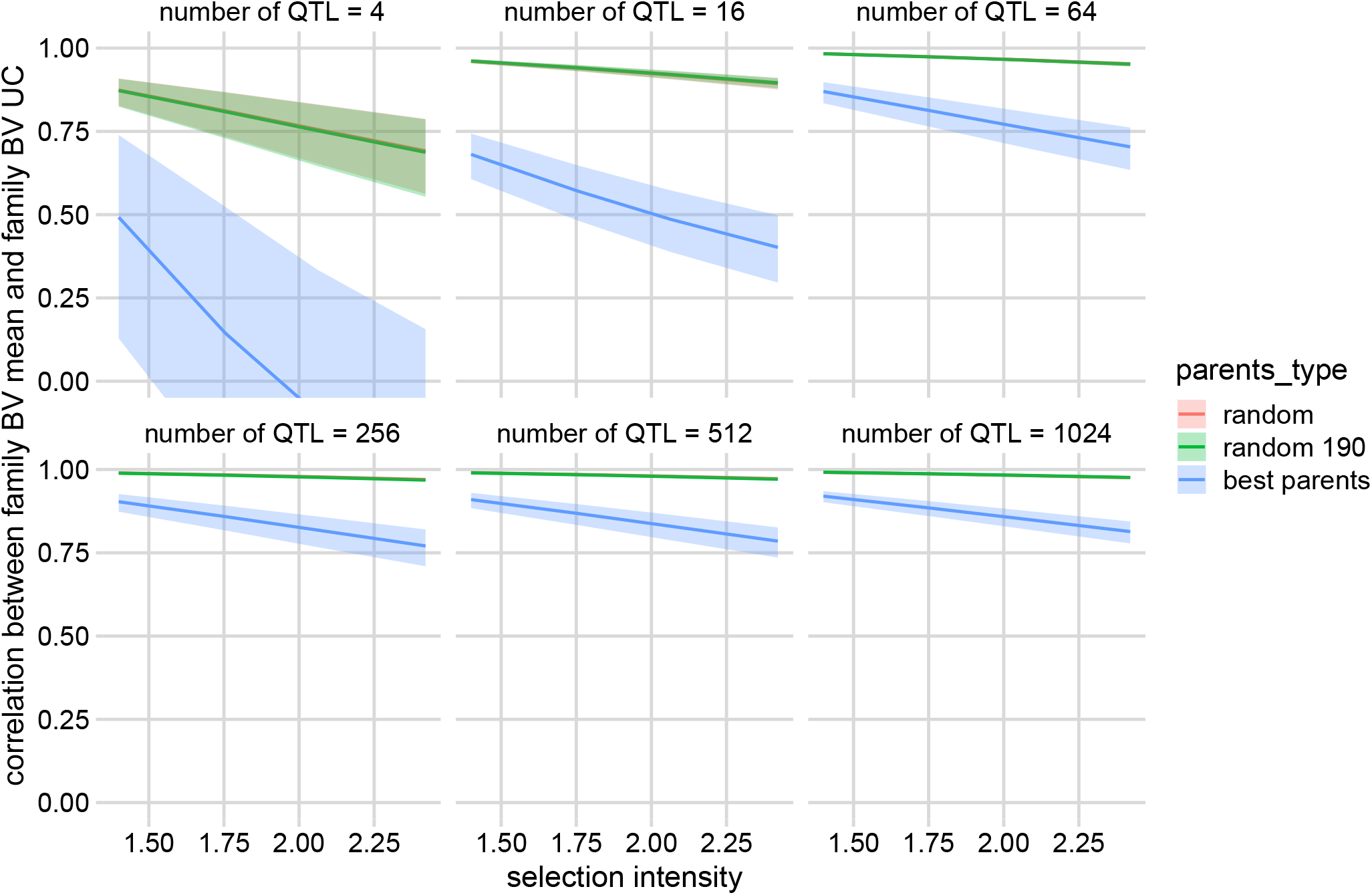
Average correlations between true family means of breeding values and true cross usefulness values across different populations of candidate crosses. Each panel shows the average correlation for simulations with a given number of QTL. Lines represent smoothed averages of the correlation between the true family mean of breeding values and the true usefulness, plotted as a function of selection intensity. The bands depict the standard errors of the correlations. Different colored lines and bands show correlations calculated from different populations of families. Red: the original 520 randomly selected families (“random”). Green: a random selection of 190 families from the 520 families (“random 190”). Blue: 190 families representing all pairwise crosses among the best 20 parents from each genetic architecture (“best parents”). The “random” and “random 190” families overlay almost perfectly.

To test whether our predictions of family standard deviations were sufficiently accurate in the “best” families to accurately predict family usefulness, we used our previously trained BayesC model to predict family standard deviations and usefulnesses. Prediction accuracies were lower than what we achieved among crosses between random parents (Figure 6, ranging from 0.2 to 0.7 vs 0.5 to 0.9, Figure 3), but increased as the number of QTL increased, particularly when the heritability was high. We observed a moderate benefit of the predicted usefulness over the predicted family mean in the high heritability scenario, but minimal benefit in other scenarios. This suggests that predictions of family standard deviation were only good enough to be useful when heritabilities were very high and even then, only if elite parents were pre-selected for families.

**Figure 6.**
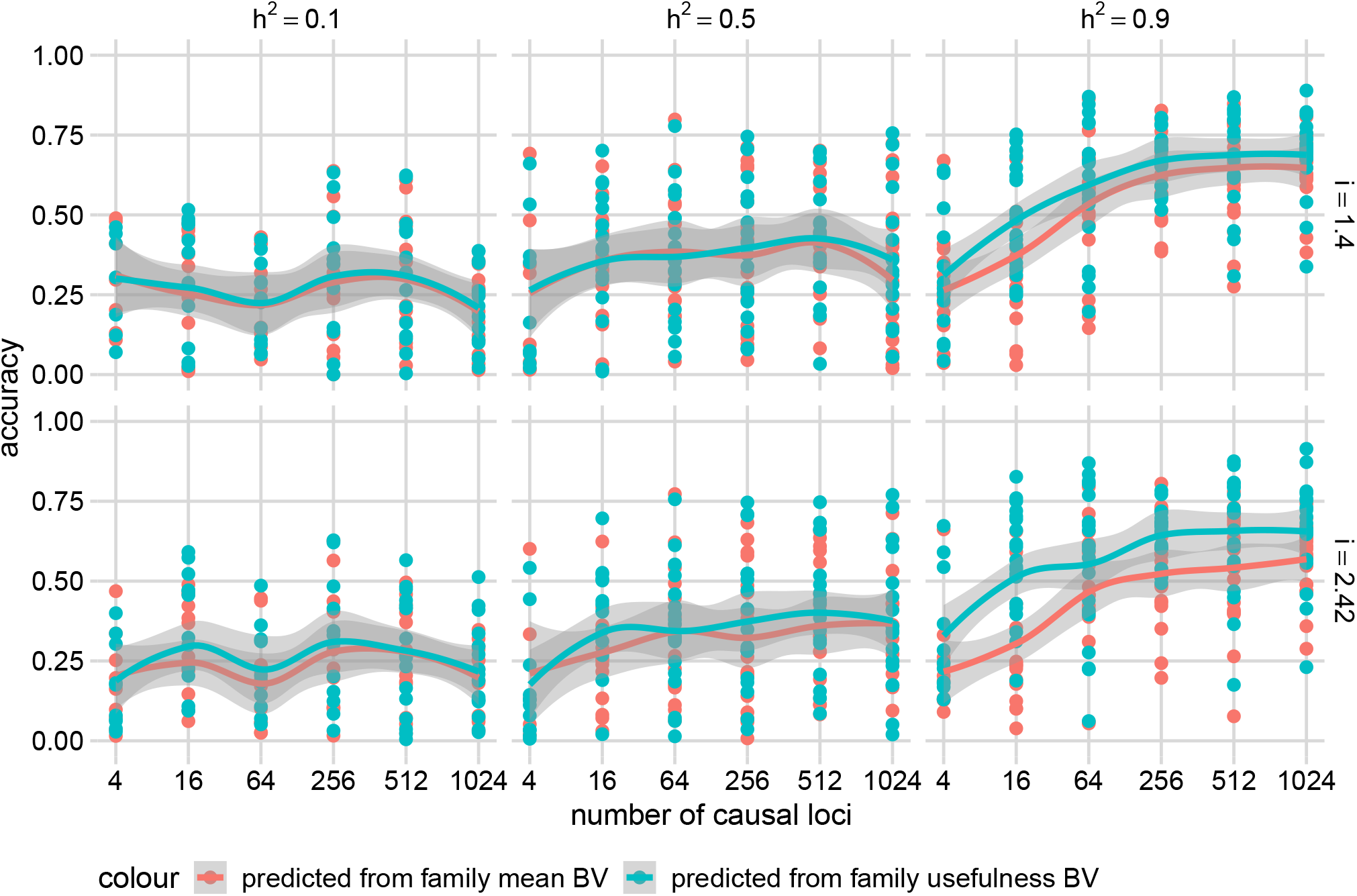
Prediction accuracies of the usefulness across the “best” families. Panels show Pearson’s correlations (accuracy, y-axis) between the true family usefulness and predicted family mean (red) or predicted family usefulness (blue) based on a BayesC genomic prediction model trained on parental phenotypes simulated with different heritabilities (*h*2) and different numbers of causal loci (x-axis), across different selection intensity (*i*), from the “best” families. 20 simulations were run at each combination of *h*2 and number of causal loci. Points show measured accuracies in each simulation across each *i*. Curves are calculated with the ‘geom_smooth()’ function and show smoothed estimates of the standard error of the mean accuracy across simulations.

Together, these results show that the true family mean is highly correlated with the true family usefulness under most scenarios, and therefore that attempting to predict the standard deviation of each family is not helpful when evaluating potential crosses, even if we could predict those standard deviations relatively accurately. This complements findings from biparental populations created from inbred parents Bernardo (2014b). However, in most realistic scenarios, the accuracy of predicting family standard deviation is not high, which would render the prediction of family standard deviation even less useful. Wolfe *et al*. (2021) mentioned that the addition of poorly predicted family standard deviation would add more noise to the family usefulness criterion, echoing our conclusion that the prediction of the family standard deviation is not helpful.

One limitation of our study is that we only simulated additive genetic architectures which is justified if we are only interested in population improvement. Under purely additive architectures, the usefulness of a cross is simply a function of the average usefulness of each parent. In this case, usefulness can be used equally as a line or mate selection metric. However, for variety release in a clonally propagated crop like strawberry, the usefulness criterion can be extended to total genetic values where dominance and epistasis effects will additionally be importantWolfe *et al*. (2021). In this case, if the within-family variance in total genetic values becomes large enough so that the variance of *iσ*_*i*_ is of similar magnitudes to *µ*_*i*_, the importance of usefulness would increase. However, predicting dominance and epistatic effects from genome-wide marker data remains very challenging and generally requires much larger sample sizes than are currently possible in most breeding programs. Wolfe *et al*. (2021) found little benefit from predicting dominance variance. In future work, we could extend our simulations to include dominance and epistatic variance to confirm these intuitions.

## Conclusions

In this study, we discuss whether selecting crosses based on predicted values of the usefulness of each cross could improve the rate of gain in breeding programs. Predicting the usefulness requires predicting the standard deviations of breeding values among the offspring produced by each cross. Our results show that predicting usefulness rarely improves outcomes relative to simply predicting the mean breeding value of each family, only in scenarios with high heritability, high selection intensity, and a small number of QTL (Figures 3 4a), and when crosses are only considered among “best parents” by the ranks of true breeding value (Figure 5). However, even in these scenarios, the relative gain in accuracy was small, suggesting that usefulness is rarely useful when making crosses in a breeding program like the UC Davis strawberry breeding program.

